# Hyperbolic geometry of gene expression

**DOI:** 10.1101/2020.08.27.270264

**Authors:** Yuansheng Zhou, Tatyana Sharpee

## Abstract

Understanding the patterns of gene expression is key to elucidating the differences between cell types and across disease conditions. The overwhelmingly large number of genes involved generally makes this problem intractable. Yet, we find that gene expression patterns in five different data datasets can all be described using a small number of variables. These variables describe differences between cells according to a hyperbolic metric. We reach this conclusion by developing methods that, starting with an initial assumption of a Euclidean geometry, can detect the presence of other geometries in the data. The Euclidean metric is used in most of current studies of gene expression, primarily because it is difficult to use other non-linear metrics in high dimensional spaces. The hyperbolic metric is much more suitable for describing data produced by a hierarchically organized network, which is relevant for many biological processes. We find that the hyperbolic effects, but not the space dimensionality, increase with the number of genes that are taken into account. The hyperbolic curvature was the smallest for mouse embryonic stem cells, stronger for mouse kidney, lung and brain cells, and reached the largest value in a set of human cells integrated from multiple sources. We show that taking into account hyperbolic geometry strongly improves the visualization of gene expression data compared to leading visualization methods. These results demonstrate the advantages of knowing the underlying geometry when analyzing high-dimensional data.

## 1. Introduction

Many processes in biology exhibit hierarchical organization, as evidenced by phylogenetic trees and clustering clades that are used to characterize cells [1], proteins [2], the activity of metabolic networks within cells [3, 4] and human brain functional networks [5]. Hiearchical organization is most congruent with a hyperbolic metric [6], and not with the Euclidean one [7]. Unfortunately, the Euclidean metric currently underlies most clustering and visualization algorithms, including k-means clustering [8], local linear embedding [9], t-distributed Stochastic Neighbor Embedding (t-SNE) [10, 11], and Uniform Manifold Approximation and Projection (UMAP) [12], etc. Therefore, there is a need to develop methods that can detect situations where hyperbolic metric is more suitable than the Euclidean one, and visualize these data with the appropriate metric. We address this need here by developing two approaches. First, we develop a non-metric multidimensional embedding (MDS) in hyperbolic space which, combined with Euclidean MDS, can quantitatively detect intrinsic geometry and characterize its properties. Second, we develop hyperbolic t-SNE to perform t-SNE embedding in a hyperbolic space instead of a Euclidean one [10]. Applying hyperbolic t-SNE to human gene expression data [13], we find that it offers more accurate embeddings than those that use Euclidean metric, including the PCA, t-SNE and UMAP [12].

## 2. Results

### 2.1. Non-metric MDS outperforms metric-MDS in geometry detection

Multi-dimensional scaling (MDS) has been widely used to embed a set of data points into a geometric space in a way that attempts to best preserve the distances between points in the original space. Metric MDS tries to make the embedding distances proportional to the input distances, while non-metric MDS preserves the ordinal values, allowing a monotonic nonlinear transformation between the distances. Both metric and non-metric MDS in high dimensional Euclidean space have been well studied during the past few decades. However, the MDS in the hyperbolic space has not been fully developed yet. Several metric MDS algorithms have been proposed recently for embedding data into hyperbolic space, offering advantages over Euclidean visualizations in terms of distance preservation, space capacity, trajectory inference and unseen data prediction [14, 15, 7, 16, 17, 18, 19, 20, 21, 22], etc. However, we find that metric MDS does not correctly distinguish between Euclidean and hyperbolic geometry of input data, but non-metric MDS does (Fig. 1). The reason is that non-metric MDS matches the ranking order instead of exact values of the data distances. The resulting nonlinear distortions in embedding distances can be used as indicators for a geometry mismatch between data and embedding points. When using non-metric MDS, we illustrate that as soon as there is a mismatch between native and embedding geometry, a nonlinear distortion appears in the scatter plots of embedding distances versus input data distances (Fig. 1A-B). These scatter plots are known as Shepard diagrams [23]. When Euclidean data is embedded into a hyperbolic space, the Shepard diagram has negative curvature (Fig. 1A). When hyperbolic data is embedded into Euclidean space, the Shepard diagram has a positive curvature (Fig. 1B). Thus, the curvature of the Shepard diagram indicates the difference in geometric properties between the embedding and native spaces, and in particular could indicate the difference in curvature of geometry. When using the metric MDS, the Shepard diagram shows increased spread (Fig. 1D) but does not yield a nonlinear relationship when embedding Euclidean data to hyperbolic space (Fig. 1C). The reason is that Euclidean distances can be embedded into the faster-expanding hyperbolic space masking the distortion of distances, and this does not happen in the non-metric MDS [23]. In what follows we apply non-metric MDS to synthetic and several real gene expression datasets to detect their hidden geometry. In what follows, we will use MDS to refer non-metric MDS for brevity.

**Fig 1:**
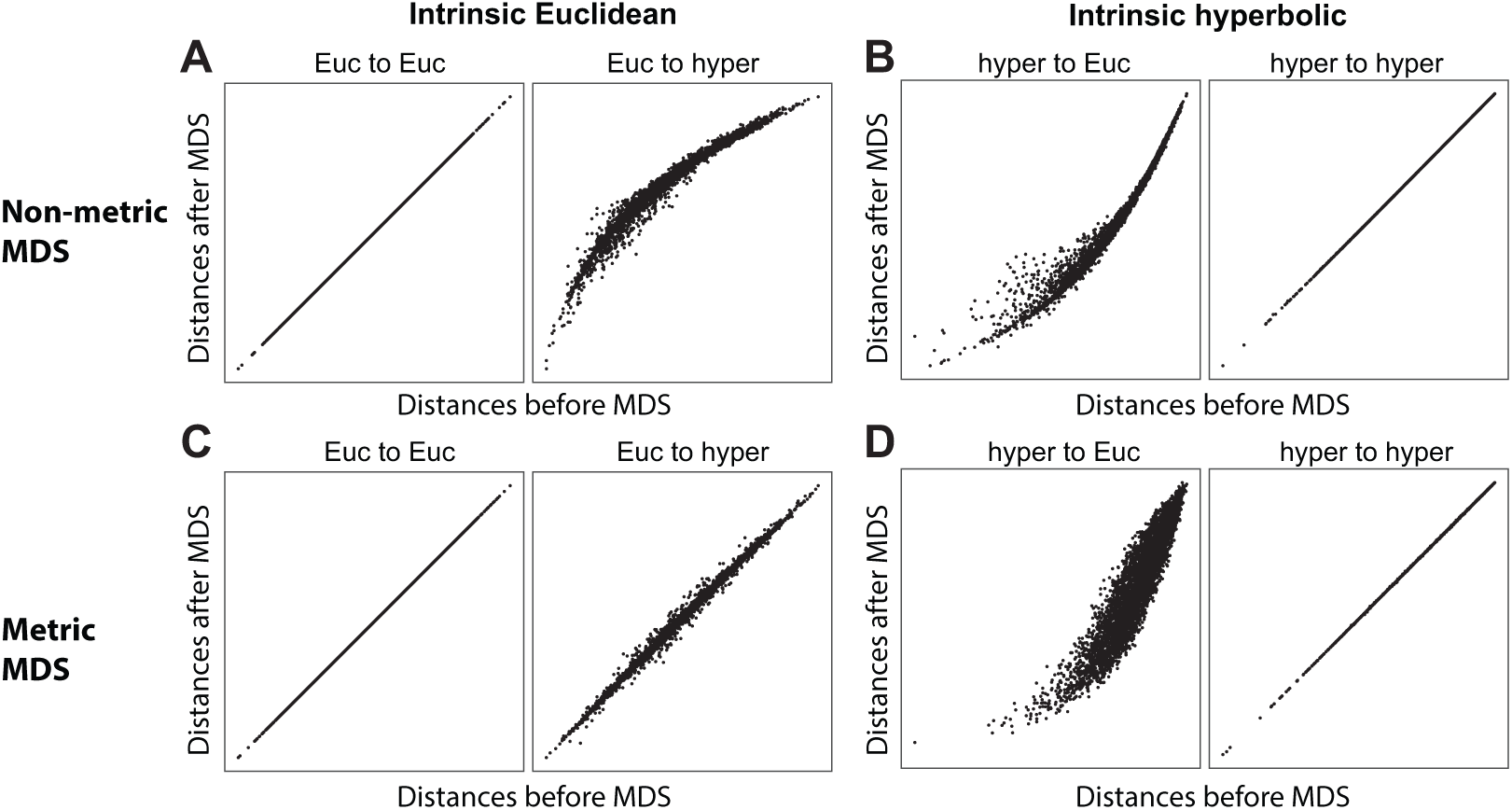
Shepard diagram for metric MDS and non-metric MDS applied to synthetic geometric data with either Euclidean or hyperbolic native geometry. These simulations were produced by randomly sampling 100 points in 5D Euclidean (A,C) and hyperbolic space (B,D), computing geometric distances using the corresponding distance metrics, and using either non-metric MDS (A-B) or metric MDS (C-D) to embed the points to 5D Euclidean and hyperbolic space. The hyperbolic radii in data sampling and hyperbolic MDS embedding are *R*_data_ = *R*_model_ = 3.0. The curvature of Shepard diagrams reflects the difference in geometry between the embedding and native spaces.

### 2.2. Synthetic geometric data

When cells are characterized according to the expression of thousands of genes, the number of genes represents the nominal dimension of the representation space. However, the real dimension of the gene expression space might be much lower. Furthermore, the true geometry of the hidden space is not necessarily Euclidean. Therefore, in this section we analyze the signatures of low dimensional hyperbolic or Euclidean geometry in the situation where each data point is described with respect to large number of variables. We focus on hyperbolic and Euclidean geometries, because hyperbolic geometry describes hierarchically organized data whereas Euclidean metric is often the only feasible geometric metric for computing distances of high dimensional vectors. In the synthetic examples below, the hidden low dimensional geometry is set to be either hyperbolic or Euclidean. To prepare for the analysis of experimental data, the high dimensional data points are evaluated using Euclidean metric, and then the Shepard diagrams are generated by applying MDS to reveal geometric properties.

First, we analyze the case where data has a 5D Euclidean underlying geometry. To simulate this case we randomly sample 100 points from a 5D Euclidean space, and use Euclidean MDS (EMDS) to embed the points to 5D, 10D, 50D and 100D space respectively (Fig. 2A, left). This step emulates the representation of real data where each data point is described by a large number of measurements (e.g. transcriptome) according to which each cell is characterized, and the distances between points are measured according to a Euclidean metric. The embeddings with different number of dimensions correspond to cases where measurements are taken with respect to different number of genes. As expected, the distances of synthetic 5D Euclidean points can be preserved without distortion in Euclidean representation spaces of higher dimensions, as evidenced by the linearity of Shepard diagrams in left column of Fig. 2A. Next, we apply EMDS (Fig. 2A, middle) and hyperbolic MDS (HMDS) (Fig. 2A, right) to the points in the Euclidean representation space, as we did in Fig. 1A-B. As one can see in Fig. 2A, Euclidean embeddings of these data do not generate distortions in the Shepard diagrams but hyperbolic embeddings yield Shepard diagrams with negative curvatures and are largely independent of embedding dimensions. This indicates that the data has an underlying Euclidean geometry.

**Fig 2:**
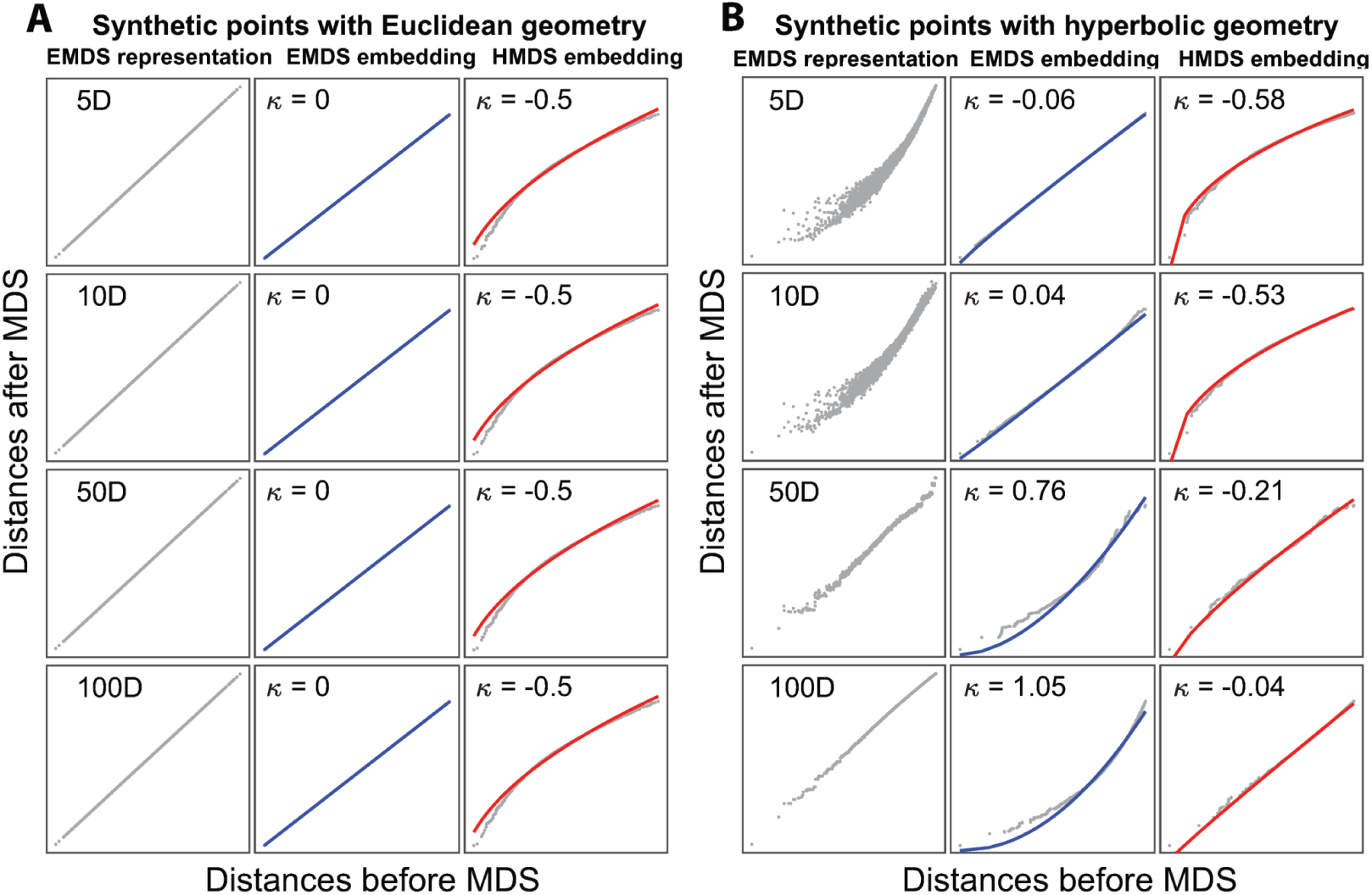
Illustration of geometry detection on synthetic data. (A) Randomly sampled 100 points from 5D Euclidean space are embedded into 5, 10, 50 and 100 dimensional Euclidean spaces (left), followed by subsequent embeddings into 5D Euclidean space (middle) or hyperbolic space (right). The solid lines represent the fits using Eq. (1), the Shepard diagram curvatures *κ* are shown at the top of each panel. (B) Same analysis for 100 sampled points from a 5D hyperbolic space with *R*_data_ = 3.0. The hyperbolic radii used in HMDS are *R*_model_ = 3.0 in both (A) and (B).

To quantitatively characterize Shepard diagrams we fit them using:

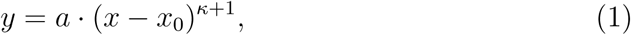

where *x* and *y* represent distances of points before and after embedding, respectively; parameter *x*_0_ = *min*(*x*) − *ϵ* is the distance offset representing the difference caused by noise from biological variations or experimental measurements, with a small *ϵ* introduced to avoid zero input values in the fitting. The key parameter *κ* characterizes the curvature of Shepard diagrams. The zero *κ* = 0 indicates pure linearity and exact match of data geometry with model geometry, while *κ* ≠ 0 means non-zero curvature and mismatch of geometry, mirroring the difference in curvature between two spaces. In the current examples, *κ* = 0 in EMDS and *κ* = −0.5 in HMDS embedding with *R*_model_ = 3, with no changes in the representation dimension (Fig. 2A). These values show signatures of intrinsic Euclidean geometry of the representation (Fig. 1A).

The situation is qualitatively different for the case where the data has hidden hyperbolic geometry, cf. Fig. 2B. Here we sample 100 points from a 5D hyperbolic space with *R*_data_ = 3.0. The initial embedding of these points into a low-dimensional Euclidean space produces distortions, indicating that using Euclidean metric to evaluate hyperbolic distances between points will not be accurate when using the same dimension of the embedding space. However, Euclidean embeddings into larger dimensional spaces can produce accurate distance representations. For example, in Fig. 2B, left column, accurate distance representation is obtained starting with ~ 50 embedding dimensions. The reason for this is that in a large dimensional Euclidean space points could be distributed along hyperbolic manifolds, approximating the true hyperbolic metric. We next apply MDS to examine the geometry of the representations by embedding them into a low dimensional Euclidean (middle column) or hyperbolic space (right column). With the increase of representation dimension, the Shepard diagram curvature *κ* increases from −0.06 (i.e. ≈ 0) to 1.05 in EMDS and increases from from −0.58 to −0.04 (≈ 0) in HMDS (Fig. 2B). These signatures (Fig. 1B) indicate that hyperbolic property is more fully preserved when points are characterized with respect to more dimensions. These analyses of model data illustrate how a combination of EMDS and HMDS can be used to elucidate the intrinsic geometry starting with the initial Euclidean representation.

### 2.3. Geometry of human gene expression data

We now apply these methods to analyze the intrinsic geometry of human gene expression from microarray data of [13]. This dataset consists of measurements of 22832 probes from 5372 human samples. The original study did principal component analysis and found that the first two principal axes described variation in biological variables corresponding to hematopoietic and malignancy properties. However, the presence and properties of the underlying geometry remain to be investigated. Several previous studies showed that gene expression was stochastic both at the single cell level and the population level [24, 25, 26], and the expression profiles of samples within the same cluster were dominated by intrinsic noise [24]. This would imply either Euclidean geometry, at least locally, or a lack of geometric structure altogether. On a global scale, biological systems usually show a hierarchical structure which would imply hyperbolic geometry [3, 5]. Therefore, we have separately probed the geometry of gene expression data at the local and global scales. To probe local geometry we apply k-means (*k* = 50) method to cluster the whole data and select 100 samples from a single cluster randomly. Similarly to Fig. 2, we use increasing subsets of genes (from 20 probes to all the 22283 probes) to represent samples and then perform EMDS and HMDS embeddings (*R*_model_ = 2.6) for geometry detection (Fig. 3A). Increasing the number of probes with respect to which samples were characterized corresponds to increasing the dimensionality of the initial Euclidean embedding as in Fig. 2. We find that this does not significantly change the Shepard diagram curvature in both EMDS and HMDS (*κ* ≈ 0 in EMDS and *κ* ≤ −0.4 in HMDS). These results match the fitting in Fig. 2A, and indicate that the samples taken from the same cluster have Euclidean structure, even when all the probes are used (Fig.3A). Additional analyses show that the Euclidean structure is indeed caused by the stochastic Gaussian expressions of genes among the samples within a cluster (Fig. S1–S2).

**Fig 3:**
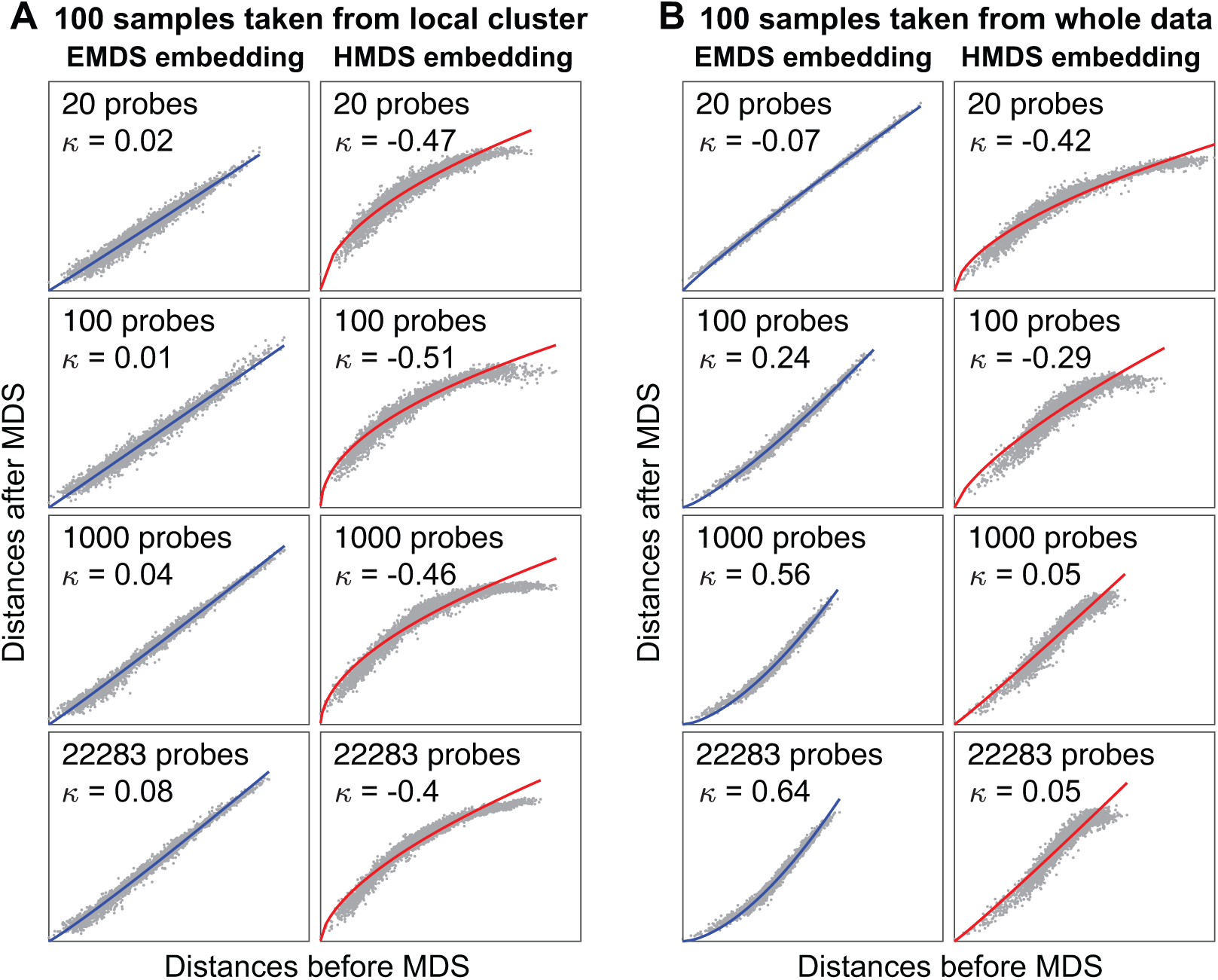
Human gene expression has locally Euclidean and globally hyperbolic hidden geometry (A) MDS embedding results for samples taken from a single k-means cluster, with distances evaluated by Euclidean metric with respect to increasing number of probes. Left and right columns show results of embeddings into 5D Euclidean and hyperbolic space, respectively. (B) Same analysis for samples taken randomly from the whole data. The hyperbolic radii used in HMDS are *R*_model_ = 2.6 in both (A) and (B).

Variations in gene expression across samples taken from different clusters, which represent different cell types, tissues and disease states, show more complicated distributions (Fig. S1–S2) and had attracted a great deal of attention [27]. To study the geometric structure of expression space globally, we select 100 samples randomly from the whole population instead of local clusters, and perform the same embeddings as in Fig. 3A. Surprisingly we find that, as the number of probes increases, the Shepard diagram curvature increases from −0.07 (≈ 0) to 0.64 in EMDS and from −0.42 to 0.05 (≈ 0) in HMDS (Fig. 3B). These fitting results match the signatures expected for hyperbolic geometry in Fig. 2B. It shows that the expression space has hyperbolic structure which becomes apparent upon including a moderately large number of genes (> 1000 probes) in the measurements.

To test robustness of this conclusion and make full use of the whole data, we repeat the sampling process 300 times both for the local sampling where samples are taken from different single clusters, and for the global sampling where samples are broadly taken from the whole data. The samples are taken with replacement. As expected, for samples taken from local clusters, the median values of curvature *κ* ≈ 0 in EMDS and *κ* < 0 in HMDS (Fig. 4A-B) even when all genes are used; these indicate Euclidean structure. For samples taken across the whole population, with increasing number of probes, the median of *κ* increases to be positive in EMDS and close to zero in HMDS (Fig. 4C-D); these signatures indicate that samples across population have hyperbolic structure when represented by a moderately large number of genes (> 1000 probes).

**Fig 4:**
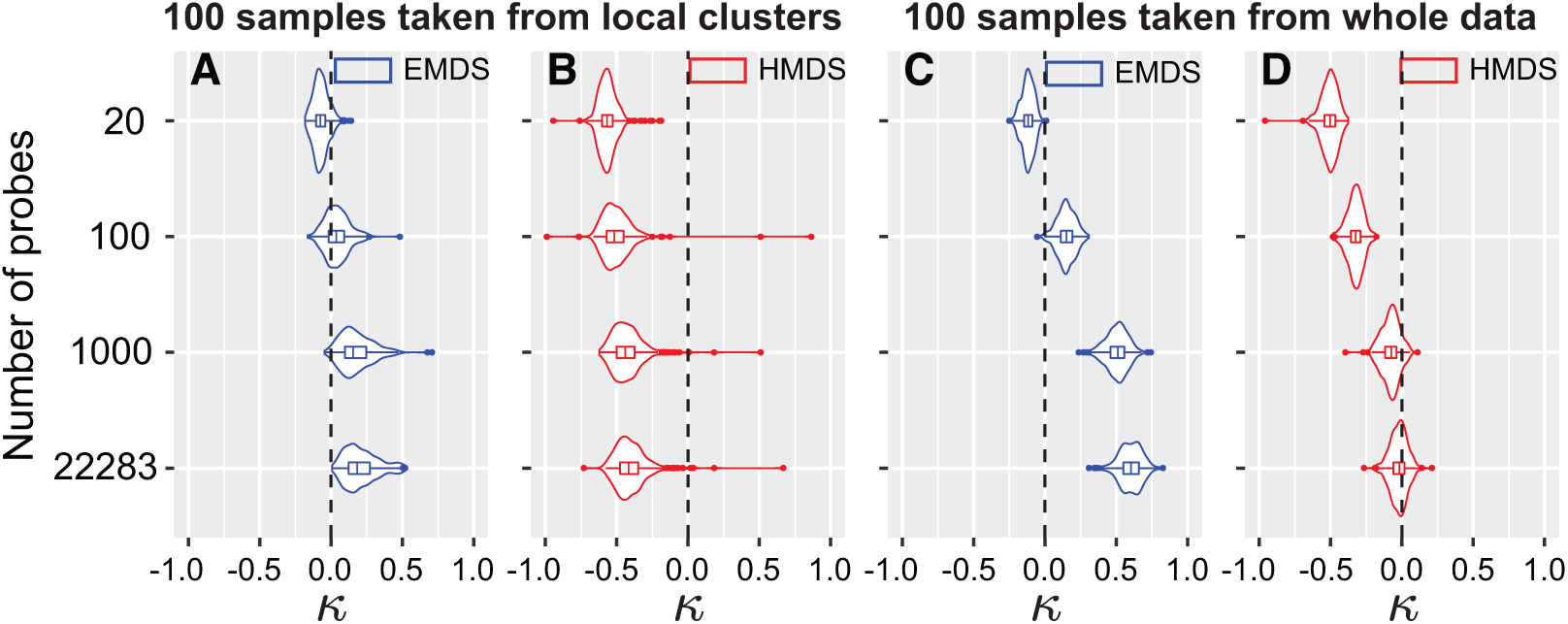
Statistics of Shepard diagram curvature *κ* across human gene expression data. (A-B) Violin plots show the curvature statistics in 5D EMDS (A) and 5D HMDS (B) across 300 repeated sampling from different k-means clusters, as a function of the number of probes. 100 data points are taken in each sampling. (C-D) Same analysis for samples taken with replacement from the whole data. The black dashed lines show *κ* = 0 and signify Euclidean geometry in EMDS (A and C) or hyperbolic geometry with *R*_data_ = *R*_model_ = 2.6 in 5D HMDS (B and D). In each plot, the width of the shape shows the probability density of different values; the central line, the left edge and right edge of the box within the shape represent the median, the 75th and 25th percentiles respectively. The line within the shape extends to the most extreme non-outlier points, the outliers are represented by dots.

### 2.4. Hyperbolic radii of embedding space for other gene expression data

In previous sections, we show that global human gene expression data can be embedded without distortion to 5D hyperbolic space with *R*_model_ = 2.6. This value was obtained by systematically screening across different *R*_model_ values to find those best matching *R*_data_ as indicated by the zero value of the curvature *κ* of the corresponding Shepard diagram fitting. In Fig. 5A we show that the curvature *κ* decreases with *R*_model_ and hits 0 at *R*_model_ ≈ 2.6, so we conclude that *R*_data_ = 2.6 in 5D hyperbolic representation. Next we examine several other gene expression datasets and determine *R*_data_ for them. We have examined single cell gene expression data from major mouse organs [28]. Applying 5D HMDS to the gene expression data from four mouse organs – brain, lung, kidney and embryonic stem cells, we find that all these data have an underlying hyperbolic structure (Fig. 5B-E). However, the hyperbolic radii of the embedding space in the case of mouse data are all smaller than those for human samples. Among the four cell types, the largest radius is found for the mouse brain cells with *R*_data_ = 1.62 (Fig. 5B), followed by mouse lung and kidney that have similar radii *R*_data_ = 1.47 and 1.45, respectively (Fig. 5B-C). Finally, the mouse embryonic stem cells have the least curved geometry with *R*_data_ = 0.98 (Fig. 5E). Because hyperbolic radius indicates the depth of the underlying hierarchical tree, these findings indicate an interesting progression with embryonic cells exhibiting the smallest degree of hierarchical organization and brain cells exhibiting the largest degree.

**Fig 5:**
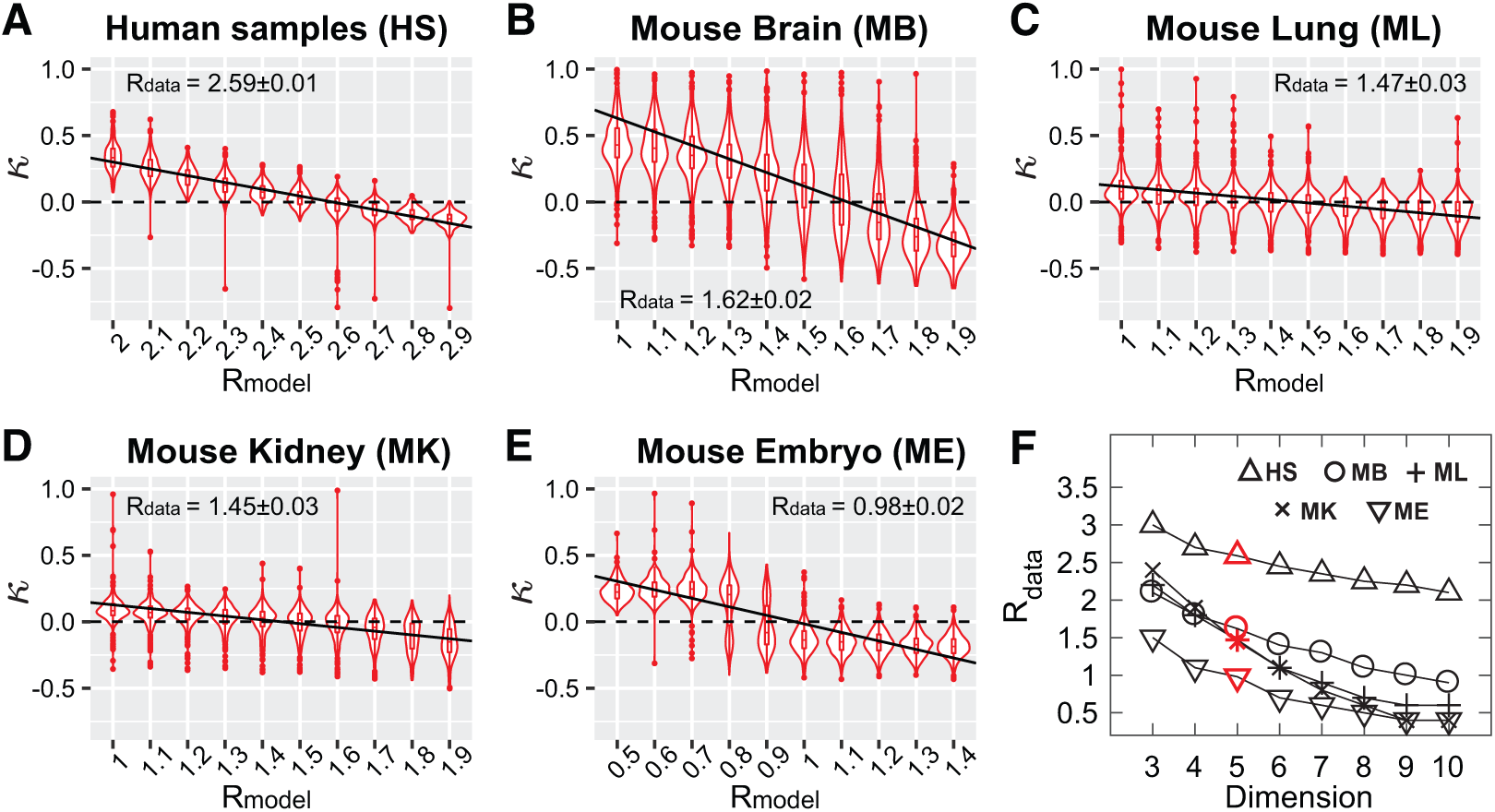
Hyperbolic radii for different gene expression data. 100 samples are taken with replacement from the whole data for 300 repeats, with all genes used. (A-E) The violin plots of Shepard diagram curvature statistics *κ* as a function of *R*_model_ for microarray of human samples in Lukk dataset (A), the microwell-seq in mouse lung (B), kidney (C), brain (D) and embryonic stem cells (E) in dataset [28]. The horizontal dashed lines show *κ* = 0, and the solid lines show the linear regression of the violin plots by *κ* = *a*_1_(*R*_model_ − *a*_2_), in which the intercept *a*_2_ is the estimate of *R*_data_. The estimated value of *R*_data_ with two times of standard deviation is shown in each panel. The HMDS embedding dimension *D* = 5 in (A-E). (F) Plots of the estimated *R*_data_ in different datasets as the function of HMDS embedding dimension.

It is also worth noting that hyperbolic radius is coupled with embedding dimension *D* by Eq. S3. Results in Figure 5A-E were obtained for a 5D hyperbolic space. In panel F, we show how *R*_data_ decreases with embedding dimensions in different data sets (Fig. 5F). We also find that minimal embedding dimension for all of these datasets is *D* = 3, and smaller dimension fails to properly embed the data.

### 2.5. Hyperbolic low dimensional visualization of human gene expression data

While MDS embedding can be used to detect intrinsic geometry, they are not ideal for low dimensional visualization. One of the primary reasons that is common to all MDS-based algorithms is that they are not designed to attract similar points together like t-SNE. Consequently, MDS-based methods achieve poor clustering results. These limitations were solved by nonlinear methods like t-SNE and UMAP, which however, are only performed in the Euclidean space. As a result, existing visualization methods may cause distortion of global structure in the data that has a global hyperbolic structure. Here we aim to adapt the t-SNE algorithm to work in hyperbolic space. To achieve this we use hyperbolic metric to evaluate global distances in the data while keeping the local clustering aspects of the algorithm. The standard t-SNE method effectively discards distance information between distant points. We recently proposed a variant of t-SNE which aims to preserve global Euclidean structure in the data [11]. The method works by adding to the similarity distance measures present in the t-SNE another term that focuses on large Euclidean distances. This produced improved and more stable global structure preservation (see Methods). Considering that human gene expression space is locally Euclidean and globally hyperbolic, we use here a hyperbolic t-SNE (h-SNE) that applies hyperbolic metric to global similarities as defined in Zhou et al. [11], while still using Euclidean metric for original local similarities. We find that this produces more accurate embeddings compared to standard methods including PCA, t-SNE, and UMAP. In particular, the Pearson correlation coefficient in the Shepard diagrams is the highest for the h-SNE method compared to the other three, with *R* = 0.767 for h-SNE compared to *R* = 0.744 for PCA, *R* = 0.430 for t-SNE and *R* = 0.627 for UMAP, cf. Fig. 6A. The distance correlation of Shepard diagram generally quantifies the quality of embedding with respect to large distances, i.e. the global inter-class structure preservation. To measure the local structure preservation, we use the silhouette score which measures the quality of clustering [29] (see Methods). Here we find that h-SNE also significantly outperforms the other three algorithms (Fig. 6B).

**Fig 6:**
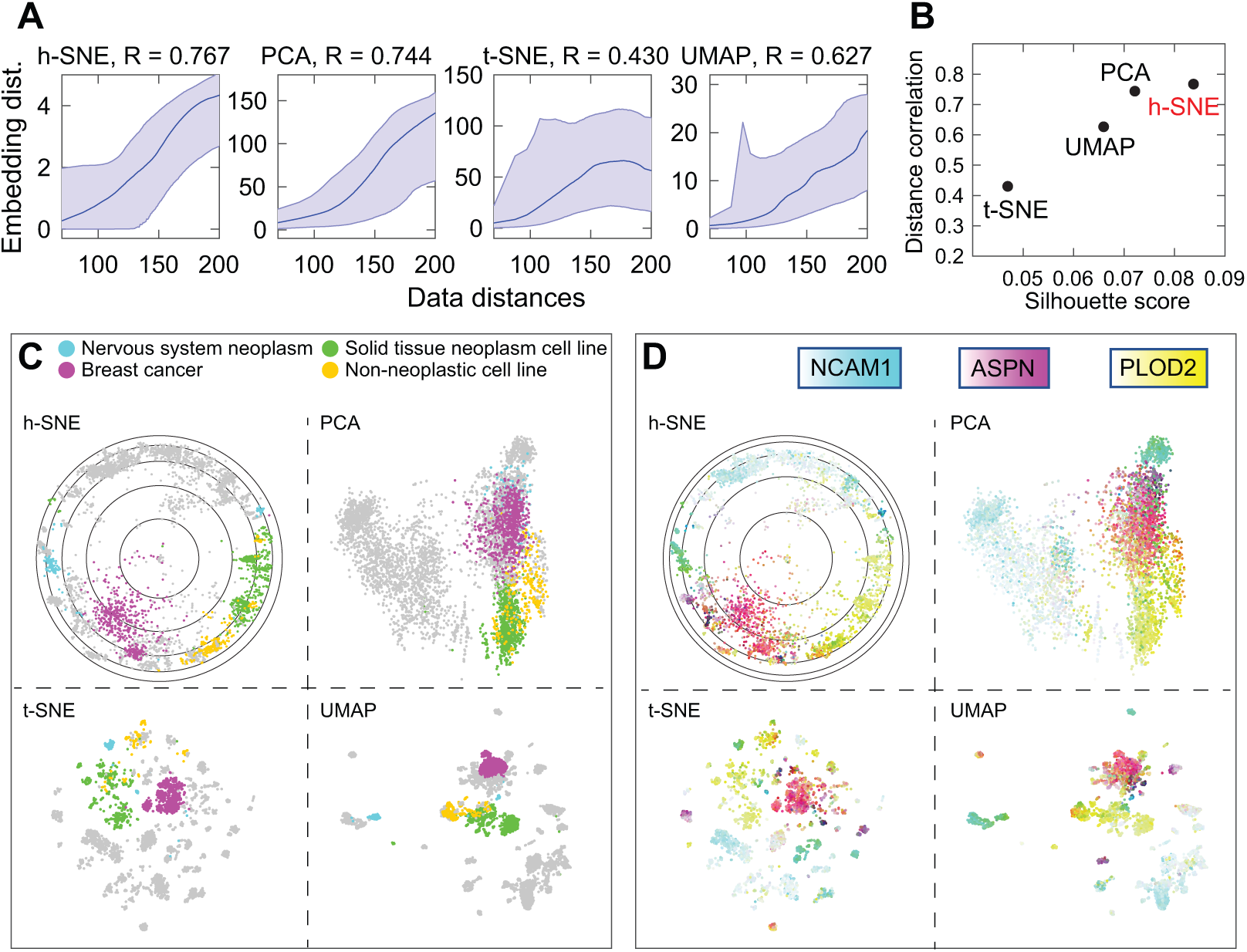
Comparison of two-dimensional visualizations of human gene expression data using h-SNE, PCA, t-SNE and UMAP. (A) Shepard diagrams of the four different mappings. The Pearson correlation coefficients of pairwise distances plots are shown at the top of each panel. (B) Quantification of local and global structure preservation using Pearson correlation coefficient of Shepard diagram (*y*-axis) and Silhouette score (*x*-axis), respectively for the four algorithms. The Silhouette score is defined as the geometric mean of the three scores obtained by using six hematopoietic labels, four malignancy labels and fifteen subtype labels respectively (see Methods). (C) In h-SNE, the data points are visualized within a 2D Poincare disk which represents a compressed version of a hyperbolic space. In PCA, t-SNE and UMAP, the points are visualized in 2D Euclidean plane. Four cell types are highlighted with color: nervous system neoplasm (cyan), breast cancer (magenta), solid tissue neoplasm cell line (green) and non-neoplastic cell line (yellow). The rest of data points are shown in gray to avoid confusion between multiple colors, see Fig. S3 for colors across all cell types. (D) The embedding samples are colored using subtractive CMY color mode according to normalized expressions of three marker genes NCAM1 (nervous system neoplasm), ASPN (breast cancer) and PLOD2 (non-neoplastic cell line).

These quantitative improvements from h-SNE are also reflected in the improved local and global visualizations that the method provides. For local visualization, the clusters identified by h-SNE are better separated with respect to 15 different tissues and disease types (Fig. S3). By comparison, the PCA representation does not separate the fifteen clusters very well, mixing nervous system neoplasm cells (cyan) with the breast cancer cells (magenta). The non-neoplastic cell line (yellow) are also not separated in the PCA representation from the solid tissue neoplasm cell line (green) (Fig. 6C, see all the 15 labels in Fig. S3). The t-SNE and UMAP methods separate clusters better but generate too many disconnected components that are difficult to be matched to sample labels (Fig. S3). In terms of global properties, the h-SNE visualization generates a clearer global organization of clusters: cells from nervous system neoplasm, breast cancer, non-neoplastic cell line and solid tissue neoplasm cell line are sequentially positioned around the disk (Fig. 6C); in addition, the two principal hematopoietic and malignancy axes can be clearly identified in h-SNE, but not in t-SNE and UMAP (Fig. S4).

The quality of h-SNE visualization is also illustrated by the topography with respect to gradient expressions of three marker genes: NCAM1 [30] for nervous system neoplasm, ASPN [31] for breast cancer, and PLOD2 [32] for non-neoplastic cell line. These marker genes are highly expressed in distinct but continuous regions in h-SNE; by comparison, the expression patterns of these three genes are more difficult to cluster in t-SNE and UMAP, or to separate in PCA (Fig. 6D).

## 3. Discussion

In this paper we developed a non-metric MDS in hyperbolic space, and showed how it can be used to detect the hidden geometry of data starting with an initial Euclidean representation. By applying this method to several gene expression datasets, we found that gene expression data exhibits Euclidean geometry locally and hyperbolic geometry globally. The radius of the hyperbolic space differed depending on the cell types. The lowest values were observed for embryonic cells and the highest values observed for brain cells in mouse data. Given that hyperbolic geometry is indicative of hierarchically organized data [33, 6], and the spanned radius represents the depth of the network hierarchy, it is perhaps intuitive that the largest value would be observed for highly differentiated and specialized brain cells and the smallest value for the embryonic cells.

The method that we used to detect the presence of hyperbolic geometry was based on non-metric MDS. One can also use methods from algebraic topology [34, 33] for this purpose, as has been recently demonstrated for metabolic networks underlying natural odor mixtures produced by plants and animals [33]. The advantage of the topological method is that it is very sensitive to changes in the underlying geometry, including its dimensionality and hyperbolic radius. However, this method is computationally intensive and does not scale well to large datasets. In contrast, the non-metric MDS method is computationally much faster. Therefore, we recommend to use it as a first step in determining whether the underlying geometry is hyperbolic or Euclidean. If hyperbolic geometry is detected, then radial position of embedding points can be used to arrange data hierarchically. We have also seen that taking into account hyperbolic geometry produces better low-dimensional visualizations, cf. Fig. 6.

Accurate representation of data across scales is a very active area of research [35, 36, 37]. Special attention is being devoted to developing visualization methods that can not only cluster data in a useful way but also preserve relative positions between clusters [12, 11]. In particular, preserving global data structure was one of the driving factors for the UMAP method [12]. Knowing the underlying geometry helps to position clusters appropriately and robustly map them across different runs in a visualization method. For example, the t-SNE method produces random positions of the clusters across different runs of the algorithm [38]. This problem can in part be alleviated by additional constraints on large distances [11]. Here we find that using a combination of a hyperbolic metric for large distances and Euclidean for local distances offers strong improvements in this respect.

What could be the origin of hyperbolic geometry at the large scale and Euclidean at small scale? First, any curved geometry, including hyperbolic, is locally flat, i.e. Euclidean. The scale at which non-Euclidean effects become important depends on the curvature of the space. From a biological perspective, the Euclidean aspects can arise from intrinsic noise in gene expression [24, 25, 26]. This noise effectively smoothes the underlying hierarchical process that generates the data. We find that hyperbolic effects can be detected by including measurements on as few as ~ 50 probes. Why do hyperbolic effects require measurements along multiple dimensions? The reason is that hyperbolic geometry is a representation of an underlying hierarchical process, which generates correlations between variables. These correlations become detectable above the noise once a sufficient number of measurements is made. As an example, one can think of leaves in a tree-like network, and how their activity becomes correlated when it is induced by turning on and off branches of the network. Intuitively, these correlations generate the outstanding branches of a hyperbola. We observe that these correlations can be detected by monitoring even a relatively small (~ 50) number of probes. This makes it possible to construct a global map of genes from partial measurements, and open new ways for combining data from different experiments.

## 4. Methods

### Metric and non-metric MDS in Euclidean model

Assume there sectionre *n* objects described by a set of measurements, the dissimilarity of the objects can be obtained by the experimental measurements of the objects. For example, the dissimilarities of two cells can be calculated by the Euclidean distances of the gene expression vectors. Metric MDS approximates the geometric distances *d*_*ij*_ to the data dissimilarities *δ*_*ij*_, while non-metric MDS approximates a monotonic transformation of dissimilarities of data. The transformed values are known as disparities 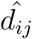. The loss function *S* in Euclidean embedding was defined as:

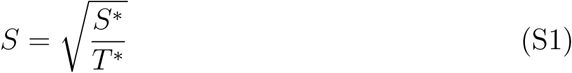

Where 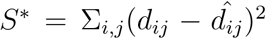, 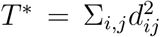. In non-metric MDS, 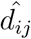 is determined using the greatest convex minorant method in Kruskal’s approach [39]. In metric MDS, disparities are equal to dissimilarities: 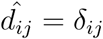.

### 4.1. Non-metric MDS in native hyperbolic model

There are many hyperbolic space representations, we will use the native representation with polar coordinates [6] in our hyperbolic MDS. The angular coordinates in the space are the same as in an Euclidean ball, the radius *R*_model_ ∈ (0, ∞) characterizes the hierarchical depth of the structure, measures the degree of hierarchy in data, and determines how points distribute in the space. The distance of two points *d*_*ij*_ is calculated as:

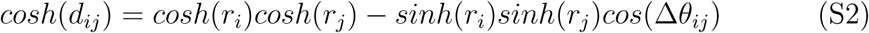

Where *r*_*i*_ and *r*_*j*_ are the radial coordinates of the two points, and ∆*θ*_*ij*_ is the angle between them. In *D*-dimensional HMDS, we initialize the embedding process by uniformly sampling points within radius *R*_model_ in the native hyperbolic model. The points directions are uniformly sampled around the high-dimensional sphere, and the radial coordinate *r* ∈ (0, *R*_model_] follows:

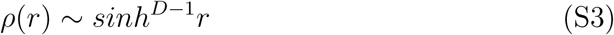

During the iteration process, we update both angular and radial coordinates according to the gradient descent of the loss function Eq.S1, and at the same time set *R*_model_ as the upper bound of the radial coordinates. The reason of setting a bound is that the coordinates in hyperbolic model are polar coordinates which cannot be normalized after each iteration as performed in Euclidean MDS, so without bound the gradient descent of loss functionsection Eq.S1 would lead to very large *r*_*i*_ and *d*_*ij*_ (since *d*_*ij*_ is in the denominator) and hence fail to preserve radial coordinates of data. By setting the upper bound for radial coordinates, the HMDS embedding can well preserve the data distances and precisely detect hyperbolic radius of data *R*_data_(Fig. S5).

### 4.2. Fitting of Shepard diagram

The Shepard diagram is linear if the geometry of input data matches the geometry of embedding space, and otherwise nonlinear. In both EMDS and HMDS, we use the power function below to fit the pairwise distances:

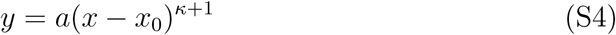

Where *x*_0_ = *min*(*x*) − *ϵ* is an offset representing the distance caused by intrinsic noise of data, a small value *ϵ* is introduced to avoid zero inputs in the fitting. The curvature *κ* describes the linearity of the fitting. *κ* = 0 indicates Euclidean input in EMDS and means *R*_data_ = *R*_model_ in HMDS. *κ* > 0 means the data is more hyperbolic than the model, and vice versa.

### 4.3. Hyperbolic t-SNE

Given a data set containing *N* data points described by *D* dimensional vectors: {**x**_1_, **x**_2_, **x**_3_, …, **x**_*N*_; **x**_*i*_ ∈ ℝ^*D*^}. The t-SNE algorithm [10] defines the similarity of two points **x**_*i*_, **x**_*j*_ as the joint probability *p*_*ij*_:

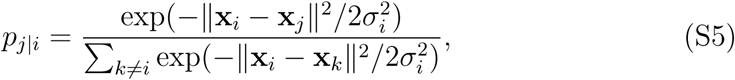

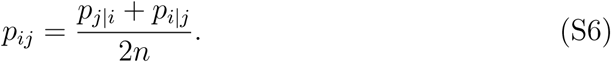

The similarity of two points **y**_*i*_, **y**_*j*_ in embedding space is defined as the joint probability *q*_*ij*_:

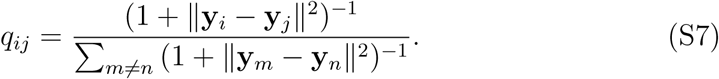

The discrepancy between the similarities of data and embedding points is the loss function, which is defined by Kullback-Leibler (KL) divergence of the joint probability *p*_*ij*_ and *q*_*ij*_:

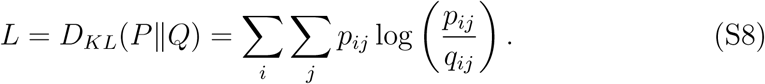

Minimizing the loss function *L* with respect to the embedding coordinates **y**_*i*_ by gradient descent gives:

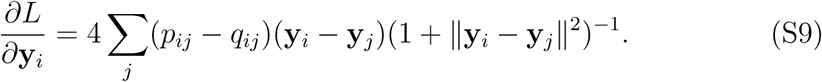

The original definitions of similarities Eqs.S5–S7 in t-SNE are sensitive to small pairwise distances among neighboring points but not to large distances between distant points. To preserve large distances, Zhou et al. [11] proposed global t-SNE algorithm that introduced global similarity terms 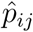 and 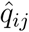 which are primarily sensitive to large distance values:

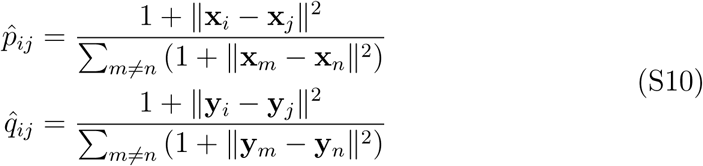

And they defined the global loss function 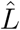 as:

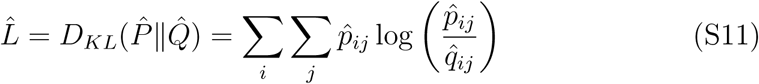

The total loss function *L*_*total*_ was then defined by combining the two loss functions using a weight parameter *λ*:

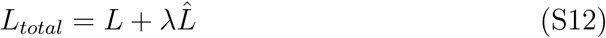

The gradient of the total loss function *L*_*total*_ gives:

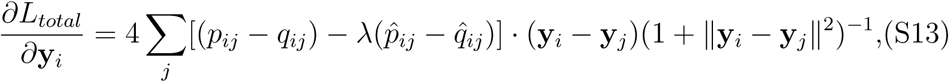

where the weight *λ* of the global loss function controls the balance between the local clustering and global organization of the data. Large *λ* values lead to more robust global distribution of clusters, but less clear classifications. Small *λ* moves back to approximate the traditional t-SNE, and will be exactly the same when *λ* = 0. In hyperbolic t-SNE, we still use native representation parametrized by *R*_model_ as in HMDS. We substitute the Euclidean distances in global similarity terms Eq.S10 by hyperbolic distances 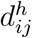 defined in Eq.S2, and change Cartesian coordinate system to polar one for all the distance calculations. Then the gradient of total loss function with respect to polar coordinates would be:

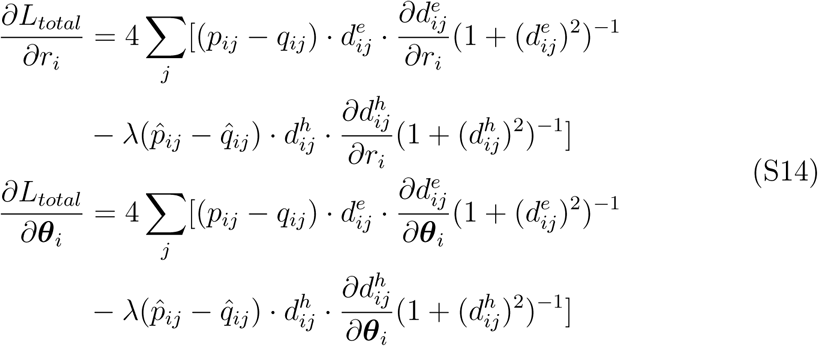

Where 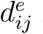 is the Euclidean pairwise distance in polar system, 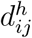 is the hyperbolic pairwise distance obtained from Eq.S2. *p*_*ij*_, *q*_*ij*_, 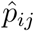 and 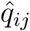 are defined by Eqs.S5–S7 and Eq.S10 with polar coordinates.

### 4.4. Parameters in visualization algorithms

In h-SNE, we set *λ* = 5 and initialize *R*_model_ = 2.7, the radial coordinates of points are not bounded during the iteration unlike in HMDS. With this parameter we run 150 repeats of h-SNE and select the embedding results with highest distance correlation of Shepard diagram. After obtaining the embedded points, we transform the points from native representation to Poincare disk model by performing the transformation on radial coordinates:

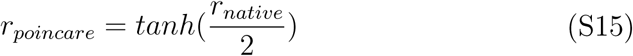

The transformed points are then displayed in Poincare disk, the circles in the Poincare disk have same intervals and the outer circle has radius which corresponds to *R* = 3.75 in native representation. The maximal radial coordinates of embedded points in optimal h-SNE is *R* = 3.18 (native representation), which is close to *R*_data_ identified in 3D HMDS (Fig. 5F). Hence we infer that the discrepancy of *R* in 2D h-SNE and 2D HMDS should not be large, despite that the latter can not be identified. In UMAP, we screen a wide range of the combination of two key parameters: number of neighbors ∈ {5, 10, 20, 50, 100} and minimal distance ∈ {0.001, 0.01, 0.1, 0.5, 0.8}, each of the 25 combinations was repeated 30 times. The optimal combination of parameters that leads to largest distance correlation of Shepard diagram is: number of neighbors = 100 and minimal distance = 0.5, and the corresponding result is shown in Fig. 6. In t-SNE we use the MATALB default parameters (Perplexity = 30) and select one with good visualization and silhouette score. In PCA, we use the first two principal components for visualization.

### 4.5. Evaluation of embedding

The Pearson correlation coefficient of Shepard diagram (embedding distances versus data distances) is used to measure the preservation of distances and global structure. For local clusters, we apply silhouette score [29] to our embedding results. Silhouette score measures the quality of data partitioning and clustering in graphical representation of objects, which in our case can be used to measure the consistency of the data configuration in 2D embeddings with the “ground truth” cluster labels. The higher score indicates better consistency with data labels. We consider all the three types of labels available – six hematopoietic properties, four malignancy properties and fifteen subtypes, and calculate the geometric mean of silhouette scores obtained by using these three labels:

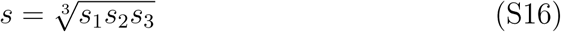

Where *s*_1_, *s*_2_, *s*_3_ represent the silhouette scores by using the three types of labeling respectively. The mean score *ϵ* is used to quantify the local structure preservation for the four visualization algorithms, which is shown in Fig. 6B.

### 4.6. Data Preprocessing and analysis

No pre-processing was done for the microarray dataset from [13]. For scRNA-seq dataset from [28], we normalized the number of reads per cell to be 1 (i.e. divide the reads array of each cell by the total number of reads in that cell), then extracted the first 1000 principal components after doing PCA. The violin plots and linear regression were performed using R version 3.6.2, the other analyses were performed using MATLAB R2017a.

## 5. Supporting information

**Fig S1:**
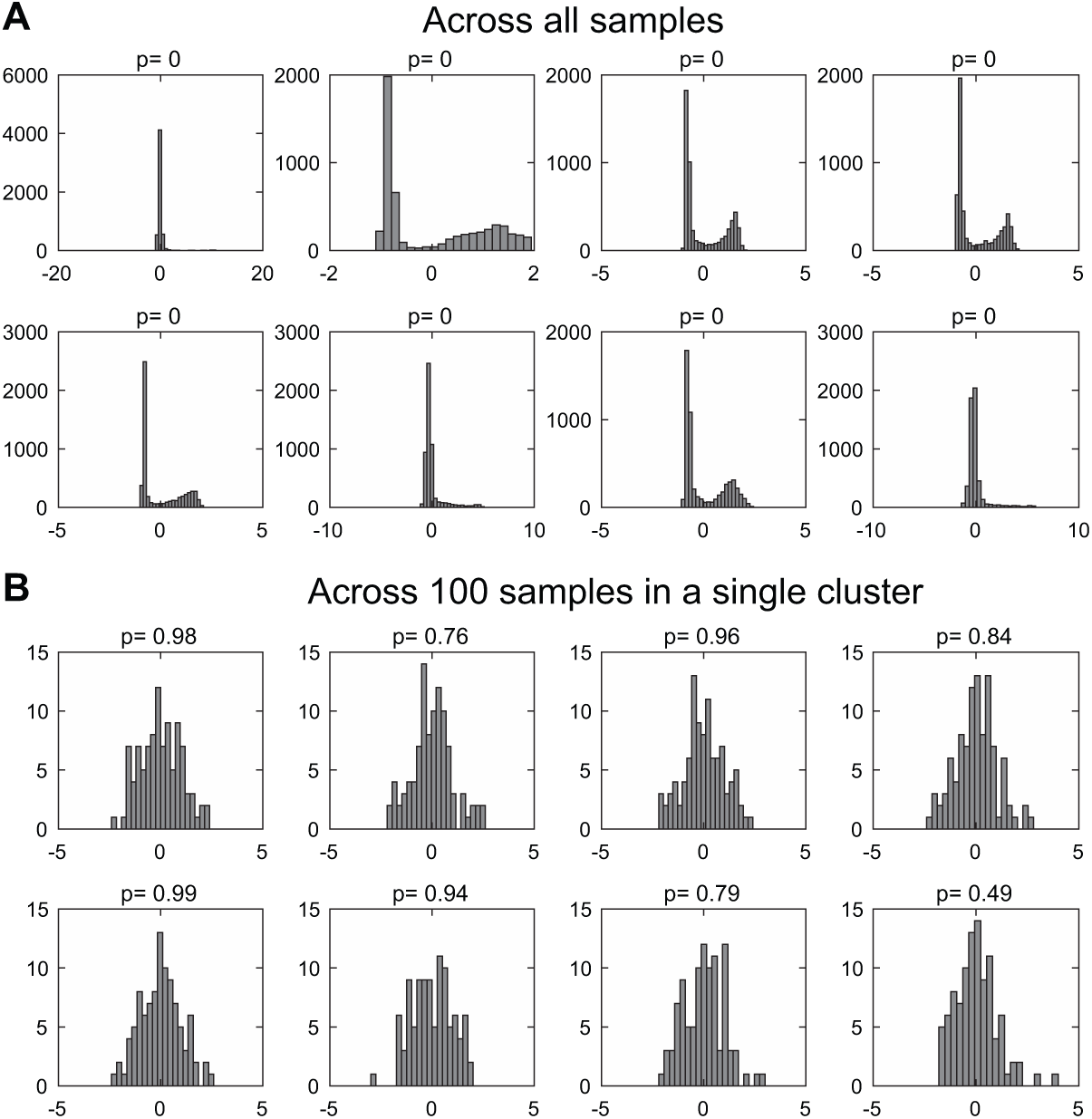
Normalized gene expression distributions across all the samples and from in a single cluster. (A) Gene expression distributions of the eight most non-Gaussian distributed probes across all the samples, p values were given by one-sample Kolmogorov-Smirnov test for Gaussianity, the null hypothesis is that the normalized distribution is standard normal distribution. (B) Gene expression distributions of the same eight probes across 100 samples in one of the k-means cluster.

**Fig S2:**
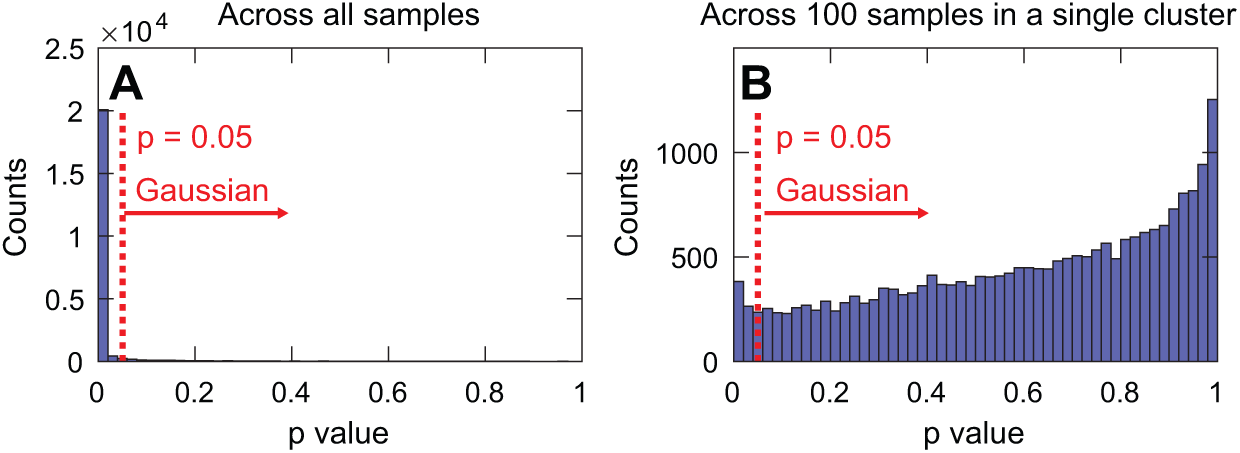
Normalized gene expression distributions follows Gaussian for samples within a cluster and non-Gaussian when samples are taken from different clusters. The p values were given by one-sample Kolmogorov-Smirnov test for Gaussianity, the null hypothesis is that the normalized distribution is standard normal distribution. (A) p value distributions of probes across the whole samples. (B) p value distributions of probes across 100 samples in one of the k-means (*k* = 50) cluster.

**Fig S3:**
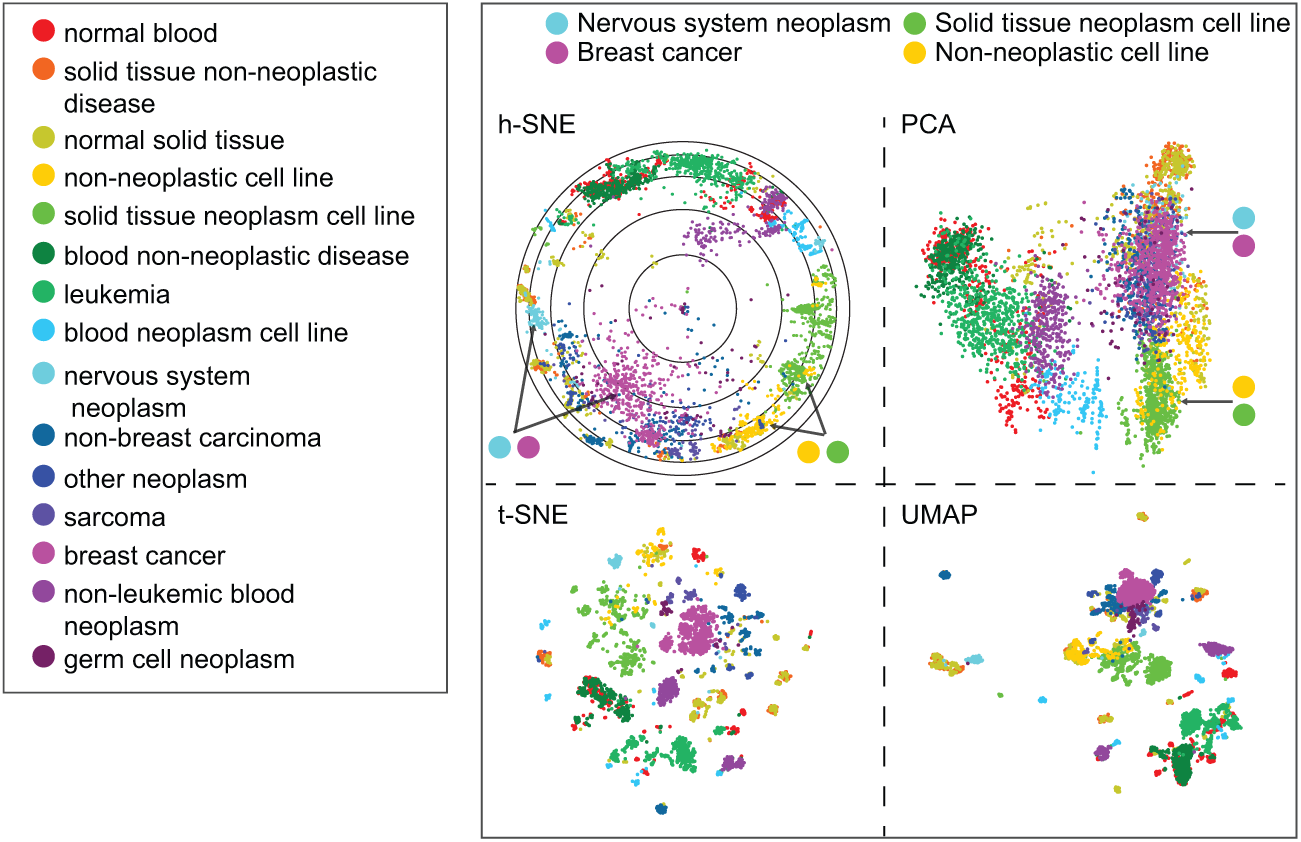
Comparison of two-dimensional visualizations of human expression data using h-SNE, PCA, t-SNE and UMAP. The samples are colored according to the 15 tissue and disease types. Other notations as in Fig. 6C.

**Fig S4:**
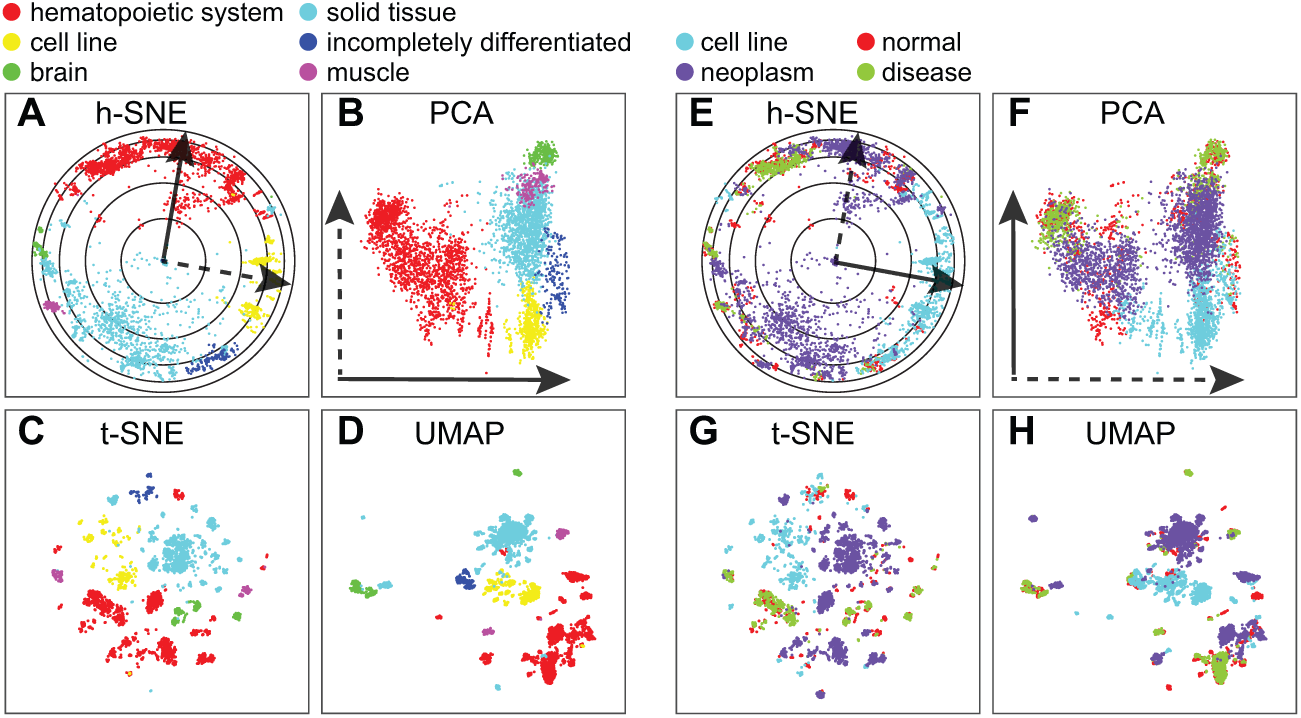
Comparison of two-dimensional visualizations of human gene expression data in different algorithms. (A-D) h-SNE, PCA, t-SNE and UMAP embeddings of human samples classified by hematopoietic properties, these labels also represent the six major clusters identified by Lukk et al. The hematopoietic axes are shown in solid lines in hSNE(A) and PCA(B) mapping. (E-H) h-SNE, PCA, t-SNE and UMAP embeddings of human samples classified by malignancy properties. The malignancy axes are shown in solid lines in h-SNE(E) and PCA(F) mapping. The six major clusters, the hematopoietic axis and the malignancy axis are hard to identify in t-SNE and UMAP mapping.

**Fig S5:**
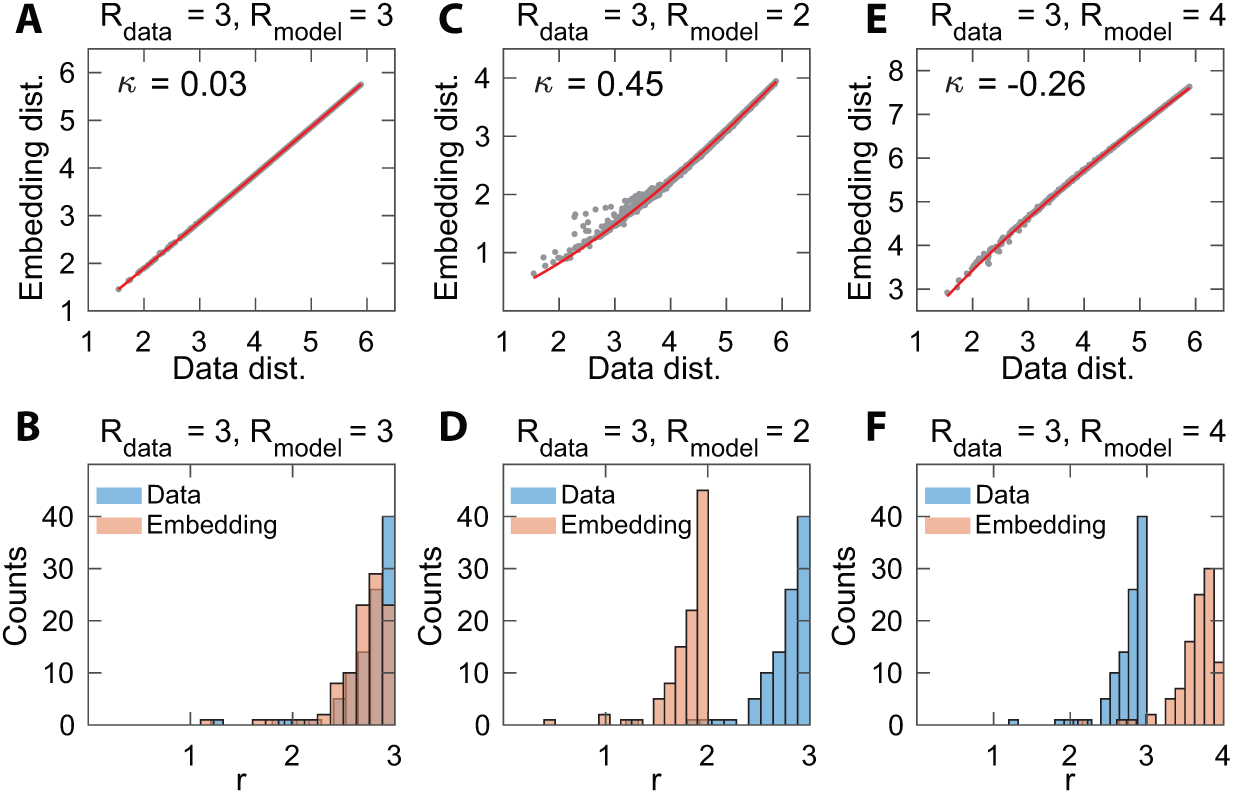
HMDS embedding of hyperbolic data with different *R*_model_. 100 points are sampled in hyperbolic space with *D* = 5, *R*_data_ = 3. The embedding dimension is *D* = 5 in (A-F). The Shepard diagram curvature *κ* is shown in panel (A,C,E). (A) Shepard diagram of HMDS embedding of the samples to 5D hyperbolic space with *R*_model_ = 3. (B) Histogram of radial coordinates *r* of 100 sample points and model points after HMDS embedding with *R*_model_ = 3. (C-D) *R*_model_ = 2. (E-F) *R*_model_ = 4.

